# Genome-wide functional analysis of hot pepper immune receptors reveals an autonomous NLR cluster in seed plants

**DOI:** 10.1101/2019.12.16.878959

**Authors:** Hye-Young Lee, Hyunggon Mang, Eun-Hye Choi, Ye-Eun Seo, Myung-Shin Kim, Soohyun Oh, Saet-Byul Kim, Doil Choi

**Affiliations:** Plant Immunity Research Center, Seoul National University, Seoul, 08826, Republic of Korea; Plant Genomics and Breeding Institute and Department of Plant Science, Seoul National University, Seoul, 08826, Republic of Korea

**Author notes:** **Corresponding Author:** Doil Choi.

**Keywords:** Plant NLR, Coiled-coil domain, Autoactivity, Singleton NLR, Cell death

## Abstract

Plants possess hundreds of intracellular immune receptors encoding nucleotide-binding domain and leucine-rich repeat (NLR) proteins. Autoactive NLRs, in some cases a specific NLR domain, induce plant cell death in the absence of pathogen infection. In this study, we identified a group of NLRs (G10) carrying autoactive coiled-coil (CC) domains in pepper (*Capsicum annuum* L. cv. CM334) by genome-wide transient expression analysis. The G10-CC-mediated cell death mimics hypersensitive response (HR) cell death triggered by interaction between NLR and effectors derived from pathogens. Sequence alignment and mutagenesis analyses revealed that the intact α1 helix of G10-CCs is critical for both G10-CC- and R gene-mediated HR cell death. The cell death induced by G10-CCs does not require known helper NLRs, suggesting G10-NLRs are novel singleton NLRs. We also found that G10-CCs localize in the plasma membrane as *Arabidopsis* singleton NLR ZAR1. Extended studies revealed that autoactive G10-CCs are well conserved in other *Solanaceae* plants, including tomato, potato, and tobacco, as well as a monocot plant, rice. Furthermore, G10-NLR is an ancient form of NLR that present in all tested seed plants (spermatophytes). Our studies not only uncover the autonomous NLR cluster in plants but also provide powerful resources for dissecting the underlying molecular mechanism of NLR-mediated cell death in plants.

## Introduction

Plants have evolved multiple immune receptors to activate defense responses against pathogen attack (1, 2). In plant cells, nucleotide-binding and leucine-rich repeat (NLR) proteins monitor pathogen invasion via direct or indirect sensing of effectors derived from pathogens (1). After recognition, activated NLRs undergo a conformational change and then initiate immune signaling (3, 4). Following NLR-mediated immune activation, a variety of defense responses is activated in infected tissues, including calcium flux, production of reactive oxygen species (ROS), activation of mitogen-activated protein kinases, biosynthesis of/ signaling by plant defense hormones, and upregulation of a subset of defense-related genes, which is often associated with the hypersensitive response (HR), a type of programmed cell death (5, 6).

Genome-wide analyses revealed that the NLR repertoire is diverse in terms of both quality and quantity among plant species, forming distinct phylogenetic clusters (7). Although the considerable effort expended over the last few decades to elucidate the mechanism of NLR-mediated resistance has enhanced our understanding of NLR classification based on function, particularly immune signaling, how multiple NLR clades function in plant immune responses remains largely unknown.

Plant NLRs were recently classified into three different categories based on mode of action: singleton, pair, and network (7). Singleton NLRs function as a single genetic unit regarding both sensing and signaling (8–10). Singleton NLRs have the potential to confer resistance when expressed in heterologous plants, even in taxonomically distant plants, as no other host factors are required for their full activity (11). In addition, most singleton NLRs possess a signaling domain that induces visibly notable cell death in heterologous plants. Paired NLRs function in pairs in which one serves as a ‘sensor’ to detect pathogens while the other functions as an immune signaling ‘executor’ (12). Ectopic expression of executor NLRs results in visible cell death without its cognate effector, which is referred to as autoactivity. The autoactivity of an executor NLR is inhibited by co-expression of paired sensor NLRs, indicating that sensor NLRs function as negative regulators of executor NLRs in the absence of pathogen infection (13, 14). Sensor NLRs in pairs have an additional integrated domain that is unusual and functions in recognizing effector proteins; this domain then relieves the regulation of the executor NLR, resulting in activation of immune responses. The typical NLR pairs are RESISTANCE TO *RALSTONIA SOLANACEARUM* 1 (RRS1)/RESISTANCE TO *PSEUDOMONAS SYRINGAE* 4 (RPS4) in *Arabidopsis thaliana* and RESISTANCE GENE ANALOG 5 (RGA5)/RGA4 in rice (*Oryza sativa*). These genes are genetically linked and oriented head-to-head so that they may share a promoter for simultaneous regulation of transcription. Although *in silico* analyses revealed that NLRs with integrated domains are common in higher plants, functional NLR pairs have been discovered in only a few plant species (15). A more complex NLR model was reported as an ‘NLR network’, primarily in Solanaceous plants. Helper NLRs, known as NLR-REQUIRED FOR CELL DEATH (NRC), are associated with phylogenetically linked sensor NLRs and exhibit functional redundancy for conferring resistance to a diverse array of pathogens (16).

NLR proteins are generally composed of three major domains: a variable N-terminus, a central nucleotide-binding (NB-ARC) domain, and a C-terminal leucine-rich repeat (LRR) domain. In general, NLR genes are divided into two major groups based on the N-terminal domain (NTD) (17, 18). One group contains the Toll/interleukin-1 receptor (TIR) domain at the N-terminus, referred as a TIR-type NLR (TNL), whereas the other possesses an NTD with a coiled-coil (CC) structure, referred to as a CNL-type NLR (CNL) (17). One group of CNLs carrying N-terminal CC domains resembling *Arabidopsis* resistance protein RPW8 was considered to represent a distinct subclass, RPW8-type CNLs (RNL), based on their function in the downstream signaling (19–21). RNLs are highly conserved in plant species, and they are necessary for the function of other NLRs (21). One particular RNL gene, N REQUIREMENT GENE 1 (NRG1), is specific to TNL-mediated immunity and partially contributes to the signaling of some CNLs (22). Other RNLs, known as ADR1s, also associate with various TNLs and function in CNL signaling (20).

Structural analyses of NLR proteins have revealed that the NTD plays multiple regulatory roles. In the case of several NLRs, the NTD interacts with host target proteins manipulated by effector proteins (23–26). Interactions between homotypic NTDs contribute to the formation of higher-order complexes of NLRs upon activation (27–29). Moreover, the NTD of NLRs is also involved in the transduction of cell death signals. Overexpression of the N-terminal region of some NLRs is sufficient to trigger cell death without cognate effector proteins (19-21, 28-31). However, how CNLs are activated and trigger cell death remains unclear.

Recent reports described the reconstitution of inactive and active complexes of an *Arabidopsis* CNL, HOPZ-ACTIVATED RESISTANCE 1 (ZAR1), with receptor-like cytoplasmic kinases (RLCKs) using structural and biochemical approaches (4, 32). Cryo-electron microscopy structural analyses demonstrated that activated ZAR1 forms a wheel-like pentamer resistosome and then undergoes a conformational change that exposes a funnel-shaped structure formed by the N-terminal α1 helix of the CC domain. These results suggest that the exposed α1 helix of the ZAR1 resistosome mediates cell death by translocating into and perturbing the integrity of the plasma membrane (PM) (32). However, whether the ZAR1 model sufficiently explains CC-induced cell death remains unclear and requires further research.

Solanaceous plants belong to a large family consisting of over 3,000 species, including a number of economically important crops, such as potato (*Solanum tuberosum*), tomato (*Solanum lycopersicum*), and pepper (*Capsicum annuum*) (33). Previous comparative analyses of NLR genes across Solanaceae genomes revealed that the NLR gene family can be classified into one TNL and 13 CNL subgroups (34). Pepper CNL-Group 10 (G10) contains 34 genes, including the known disease-resistance (R) genes *Pvr4* and *Tsw*, the products of which confer resistance to potyviruses such as *Potato virus* Y and *Tomato spotted wilt virus* via recognition of a viral effector, respectively (35, 36). Structural domain analyses revealed that the CC domain of Pvr4 is sufficient to activate cell death in the absence of the viral effector NIb (37).

In this study, we conducted a genome-wide screening in *Nicotiana benthamiana* and report here that the CC domains of G10-NLRs (hereafter G10-CCs) in pepper specifically induce HR-like cell death. Autoactive G10-NLRs and G10-CC-mediated cell death were associated with immune responses. G10-CC domains were found to localize in the PM, where the α helix of these domains plays a critical role in mediating cell death. Cell death induced by autoactive G10-NLRs and G10-CCs was not compromised in *NRC*- or *NRG1*/*ADR1*-silenced plants, suggesting that helper NLRs are not required for signaling associated with G10-NLR-mediated cell death. Surprisingly, G10-NLRs were found to be well conserved in seed plants. G10-CCs in other Solanaceae plants, such as tomato, potato, and tobacco, as well as a monocot plant, rice, induced cell death in *N. benthamiana.* We propose that G10-NLRs represent a novel singleton NLR cluster and that G10-CCs could serve as powerful resources for elucidating the underlying molecular mechanism of NLR-induced plant cell death.

## Results

### The CC domains of pepper G10-NLRs induce cell death in *N. benthamiana*

In a previous study, intact pepper NLRs were identified and assigned into 14 groups, consisting of 1 TNL and 13 CNLs (34). To identify autoactive NLRs on a genome-wide scale, genomic fragments of 436 intact pepper NLRs were cloned into the binary vector pCAMBIA2300-LIC (35, 36). The sequences of all inserts were confirmed by DNA sequencing analysis (Dataset S1), and the constructs were transiently expressed in *N. benthamiana* leaves via agroinfiltration. Among the 436 tested NLRs, only 15 (3.4%) triggered cell death (Table 1). These NLRs were randomly distributed into six groups: TNL, CNL-G5, CNL-G9, CNL-G10, CNL-G11, and CNL-NG. To determine whether these genes function as executor NLRs with their paired sensor NLRs, we examined the upstream flanking genes of the autoactive NLRs using pepper chromosome pseudomolecules, as known NLR pairs are genetically linked in a head-to-head orientation (38, 39). Although 9 of 15 autoactive NLRs had an NLR-type gene in the upstream flanking region, none were arranged in a head-to-head orientation (Table S1). Furthermore, the sequences of these upstream NLR genes did not encode an integrated decoy domain, which is a typical feature of sensors of paired NLRs. These results suggest that the conventional paired NLR immune receptors, such as *Arabidopsis* RPS4/RRS1 and rice RGA4/RGA5, probably do not exist in pepper.

Previous studies reported that the N-terminal TIR or CC domains of a specific set of resistance proteins, mostly potential singleton NLRs, induce cell death when overexpressed alone in plants (21, 40, 41). These results suggest that NLR proteins normally trigger immune signaling via the NTD. To identify NLRs carrying an autoactive NTD in pepper, a representative subset of NTDs, including those of autoactive NLRs, were transiently overexpressed in *N. benthamiana* in the same manner as the full-length NLR. The NTDs for each group were randomly chosen and represented more than 15% of all assigned NLRs from a subclade in a phylogenetic tree analysis (Table 1, Dataset S2). The NTD fragments were determined from the N-terminus up to the predicted P-loop in the NB-ARC domain. Of the 15 autoactive full-length NLRs examined, only 2 TIR domains and 3 CC domains from autoactive TNLs and G10-NLRs, respectively, induced visible cell death when expressed in *N. benthamiana* (*SI Appendix,* Table S2). Within the CNL-NG group, CC domains from pepper homologs of NbNRG1 and NbADR1 triggered cell death (*SI Appendix,* Table S2). In particular, compared with other groups, a significantly higher proportion of the CC domains of pepper CNL-G10 that were tested (70.8%) triggered cell death (Table 1 and *SI Appendix,* Fig. S1). G10-NLRs formed a distinct cluster in the phylogenetic tree that included several cloned functional R genes: *Pvr4* and *Tsw* in pepper and *RPS2* and *RPS5* in *Arabidopsis* (34). Although G10-NLRs are conventional CNLs, G10 is phylogenetically distinct from other CNL clades (34). Taken together, these data suggest that G10-NLRs have features specific to other CNLs and they may play unique a role(s) in plant immunity.

### CC-induced cell death mimics R gene-mediated HR

We next investigated whether cell death induced by autoactive G10-NLRs or G10-CCs plays a role in R gene-mediated defense responses against pathogens. An autoactive NLR, CC309, a non-autoactive NLR, CC10-1, and an autoactive full-length NLR, NLR620, were examined in this assay. A known G10-NLR gene, *Pvr4*, and its cognate Avr gene, *NIb*, of *Pepper mottle virus*, were also included as positive controls for R gene-mediated defense responses (36). Even though they share high identity (95.5% and 89.9% at the nucleotide and amino acid levels, respectively), CC309 and CC10-1 exhibited opposing phenotypes (Fig. 1A). Transient overexpression of CC309 and NLR620, but not CC10-1, triggered visible death in *N. benthamiana*. All proteins were detected in immunoblotting analysis, but the expression level of CC10-1 was much lower than that of CC309 (Fig. 1B). To exclude the possibility that the non-functionality of CC10-1 is due to low protein accumulation, green fluorescent protein (GFP) was fused to CC309 and CC10-1 at the C-terminus. Although CC309-GFP and CC10-1-GFP accumulated to similar levels, CC10-1-GFP still did not trigger cell death, indicating that the non-autoactivity of CC10-1 cannot be attributed to low expression levels (*SI Appendix,* Fig. S2).

**Figure 1.**
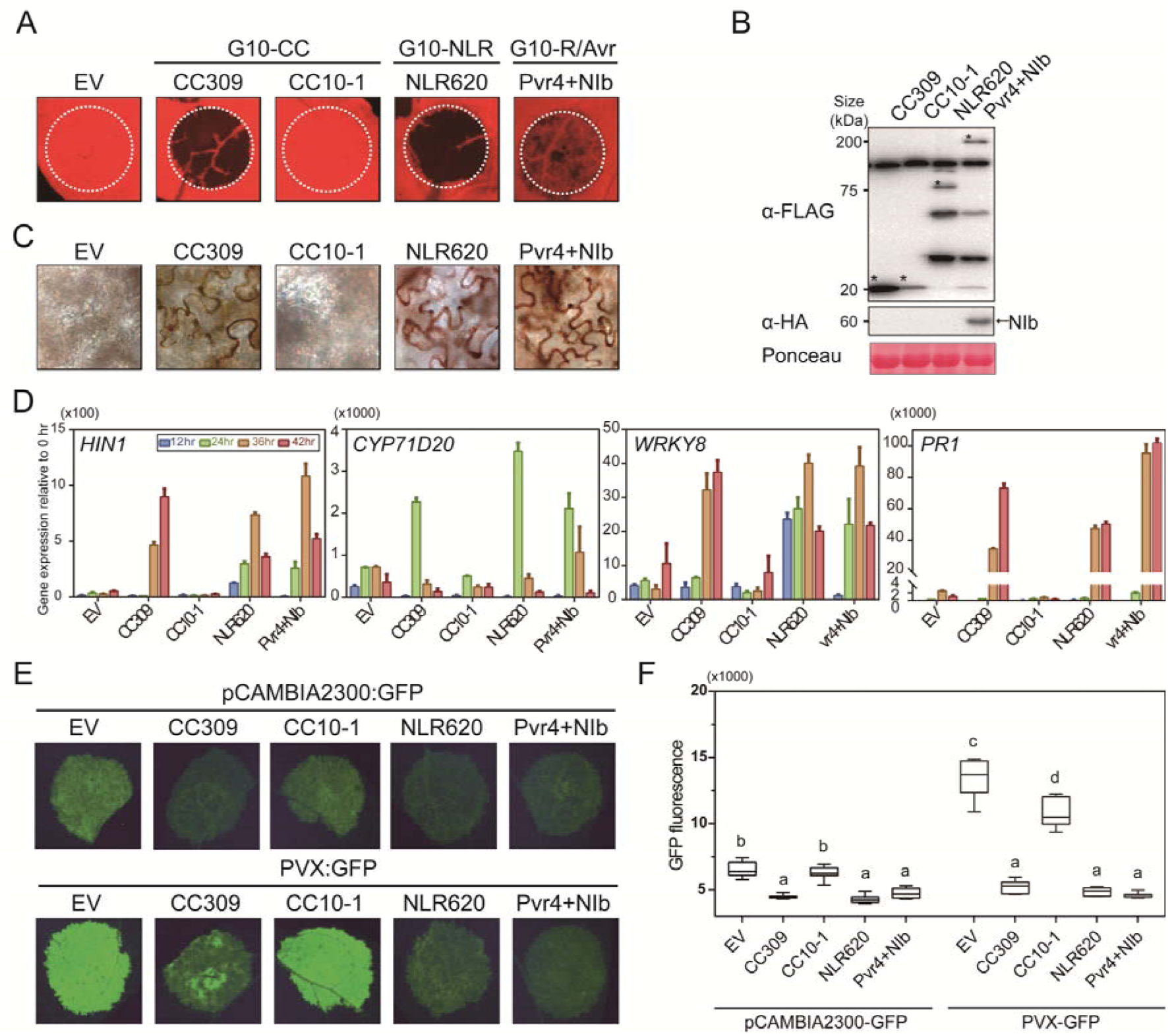
Autoactive G10-NLR and G10-CC trigger defense-related cell death. (A) CC309, NLR620 and Pvr4 with its cognate effector NIb induce cell death, but CC10-1 does not. EV (empty vector) was used as a negative control. The leaves of 4-week-old *N. benthamiana* were infiltrated with agrobacteria harboring each construct. Images were taken 3 days post infiltration. (B) Protein accumulation was confirmed by immunoblot analysis. Equal protein loading was confirmed by staining membranes with Ponceau S. Asterisks indicate the expected protein bands. (C) H_2_O_2_ accumulated in CC309-, NLR620-, and Pvr4/NIb-infiltrated leaves but not CC10-1- or EV-infiltrated leaves. Leaves were stained with DAB at 2 dpi; n = 3. (D) Expression of defense-related genes, *HIN1*, *CYP71D20*, *WRKY8* and *PR1*, was dramatically elevated by CC309, NLR620, and Pvr4/NIb. Four-week-old *N. benthamiana* leaves infiltrated with EV, CC309, CC10-1, NLR620, and Pvr4/NIb were collected at various time points. The data are shown as mean ± SD (n = 3). (E) and (F) Enhanced resistance to *Potato virus X* mediated by CC309, NLR620 and Pvr4/NIb. pCAMBIA2300:GFP or PVX:GFP was co-infiltrated with agrobacteria carrying EV, CC309, CC10-1, NLR620, or Pvr4/NIb. (F) Accumulation of GFP was visualized at 30 hpi. GFP fluorescence emission was quantified using a FluorCam with a GFP filter. Letters indicate statistical differences as determined by one-way ANOVA with Sidak’s multiple comparisons test. The above experiments were repeated two times with eight biological replicates.

Two days after agroinfiltration, we assessed hydrogen peroxide (H_2_O_2_) accumulation in the leaves using 3,3’-diaminobenzidine hydrochloride (DAB) staining. H_2_O_2_ production was observed during cell death mediated by activated Pvr4 (Fig. 1C). As demonstrated by Pvr4, cells expressing CC309 and NLR620, but not CC10-1, exhibited accumulation of H_2_O_2_. These results indicate that accumulation of ROS, which is a defense response against pathogen infection, is accompanied by cell death induced by autoactive G10-NLR and G10-CC.

To verify that ectopic expression of autoactive G10-NLR and G10-CC mimics the activation of defense-related genes, CC309- and NLR620-mediated accumulation of transcripts for four defense-related genes was monitored during the cell death process at various time points after agroinfiltration (Fig. 1D). Overexpression of CC309 and NLR620, but not CC10-1, resulted in significantly higher transcription of several genes, including the HR cell death marker gene *Harpin-induced 1* (*Hin1*), a transcription factor for physiological substrates of mitogen-activated protein kinases, *WRKY8*, a cytochrome P450 involved in sesquiterpene phytoalexin biosynthesis, *CYP71D20*, and *Pathogenesis-related gene 1* (*PR1*) (42–45). These results indicate that cell death induced by autoactive G10-NLR and G10-CC is also correlated with the accumulation of defense-related gene transcripts.

Although ROS production and upregulation of defense-related genes are typical immune responses, we cannot exclude the possibility that they also occur as a consequence of cell death. To verify that autoactive G10-NLR and G10-CC indeed trigger defense responses, we performed a co-expression assay using a *Potato virus X* (PVX) vector encoding GFP (37). We hypothesized that if autoactive G10-CC- or G10-NLR-mediated cell death is associated with activation of defense responses, replication of PVX would be suppressed in co-expressing cells. To avoid the possibility that cell death would affect the replication of PVX, the GFP signal was monitored before onset of the severe cell death phenotype (*SI Appendix,* Fig. S3). Compared with the empty vector, Pvr4 with NIb significantly reduced the abundance of PVX:GFP. Moreover, the GFP accumulation was also almost completely inhibited by expression of CC309 and NLR620 (Fig. 1E, 1F). Taken together, cell death induced by autoactive G10-NLR and G10-CC functionally mimics R gene-mediated HR cell death.

### The N-terminal **α**1 helix is critical for autoactivity of G10-CCs

We performed an alignment analysis of pepper G10-CCs to identify the conserved motif necessary for cell death activity. However, no conserved motif was found in any of the G10-CCs examined (*SI Appendix,* Fig. S4). A phylogenetic analysis based on amino acid sequences of pepper G10-CCs revealed that they are divided into three subclades, with 100% bootstrap confidence level (Fig. 2A). In the phylogenetic analysis, we found that most of non-autoactive G10-CCs (shown by open circle) belong to subclade I (Fig. 2A left panel). It was recently reported that the N-terminal α1 helix of the ZAR1 CC domain is crucial for its cell death-inducing activity based on a crystal structure model (32). This prompted us to perform further analysis of the α1 helix of subclade I members. Prediction of secondary structures using JPRED4 (http://www.compbio.dundee.ac.uk/jpred4) revealed that non-autoactive G10-CCs (CC292, CC310, CC368, and CC430) in subclade I have a shorter α1 helix than autoactive G10-CCs (Fig. 2A, right panel), and partial deletion of the α1 helix was found in non-autoactive G10-CCs (Fig. 2B). Although the α1 helix of non-autoactive CC474 is of relatively sufficient length, a deletion was found in the linker region between the α1 and α2 helixes (Fig. 2A right panel and 2B), suggesting that the deletion may disrupt formation of the structure necessary for activity. Taken together, these results suggest that the cell death-inducing activity of non-autoactive G10-CCs in subclade I is associated with the deletion in the α1 helix.

**Figure 2.**
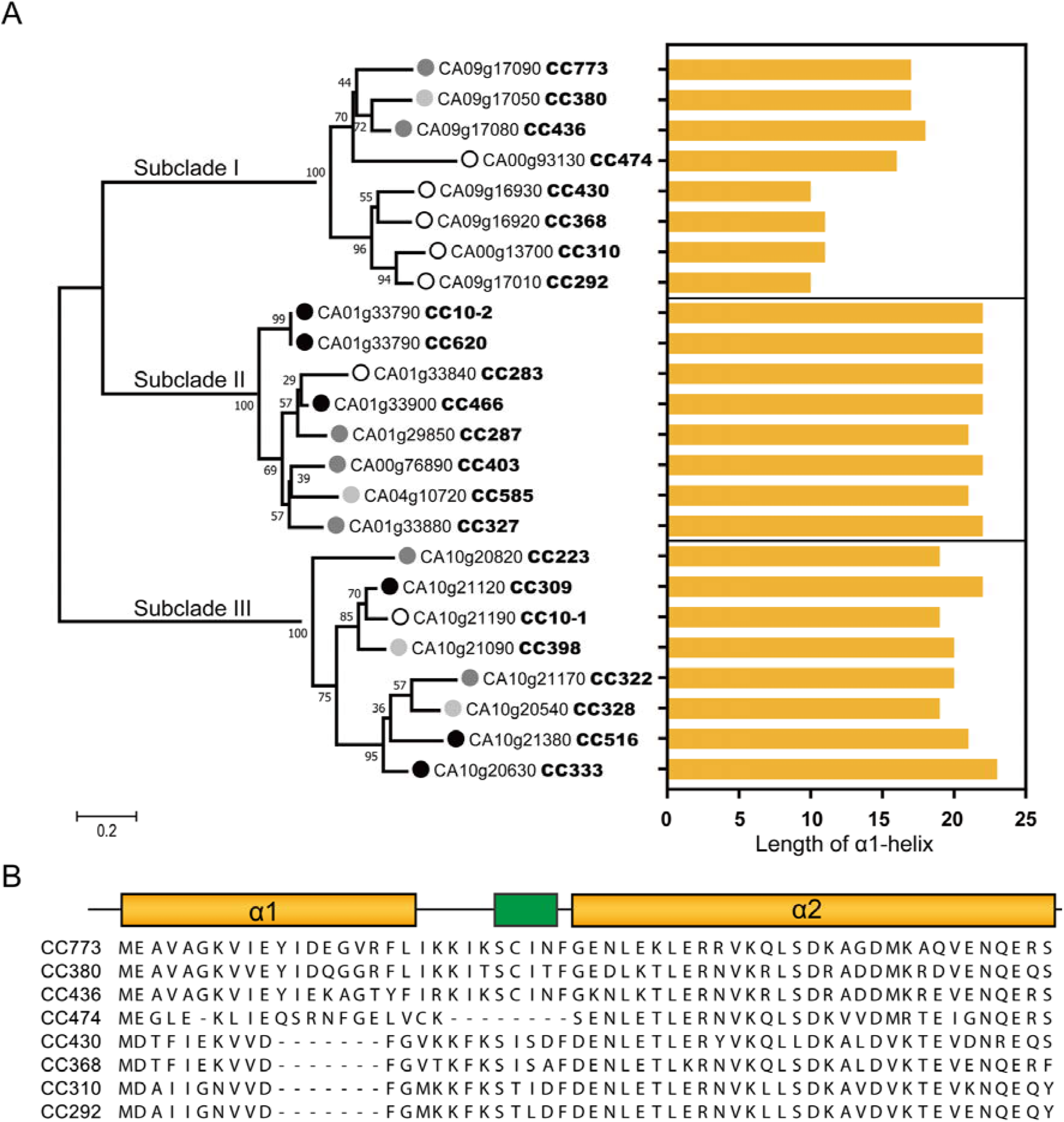
The first α-helix region is critical for autoactivity of G10-CC domains. (A) Phylogenetic tree of G10-CCs (left) and length of the first α-helix (right). The phylogenetic tree was constructed based on amino acid sequences of G10-CC domains. The maximum-likelihood model was used, and bootstrap analysis was performed with 1,000 replicates. The degree of cell death is represented as a grayscale circle to the left of the gene IDs (closed circle, strong; dark gray circle, medium; light gray circle, weak cell death; open circle, no cell death). (B) Alignment of CC domains in subclade I and schematic representation of the secondary structure of subclade I CC domains. Non-autoactive CC domains (CC474, CC292, CC310, CC368, and CC430) are deficient in an intact first α-helix. The α-helix and β-sheet are represented as yellow and green boxes, respectively.

Interestingly, CC10-1 of subclade III is the only non-autoactive CC domain that has an α1 helix without deletion. Sequence alignment with CC309 revealed a difference in the 12-aa N-terminal region (Fig. 3A, *SI Appendix,* Fig. S4). The N-terminus of the CC domain of NLRs has been reported important for both cell death activity and subcellular localization of the NLR protein (32, 46). In the case of *Arabidopsis RPS5*, a known resistance gene in G10, N-terminal acylation–mediated PM localization is essential for cell death activity, although the RPS5 CC domain also requires the NB-ARC domain to trigger cell death (46). However, pepper G10-CCs do not contain the predicted S-acylation site in the N-terminus. A transmembrane region spanning residues 1–30 was predicted by TMHMM Server ver. 2.0 with low probability in CC309, but not CC10-1 (*SI Appendix,* Fig. S5). We hypothesized that the sequence variation in the α1 helix between CC309 and CC10-1 has an effect on subcellular distribution and consequently results in the observed difference in activity. We examined the subcellular distribution of G10-CCs-GFP in *N. benthamiana* using confocal microscopy. GFP signals associated with CC309-GFP and CC10-1 were detected at the PM and colocalized with FM4-64, a fluorescent dye specific for the PM (Fig. 3B). Moreover, other non-autoactive G10-CCs in subclades I and II also localized at the PM (*SI Appendix,* Fig. S6). These results suggest that non-autoactivity of G10-CCs is not due to mislocalization to other cellular compartments.

**Figure 3.**
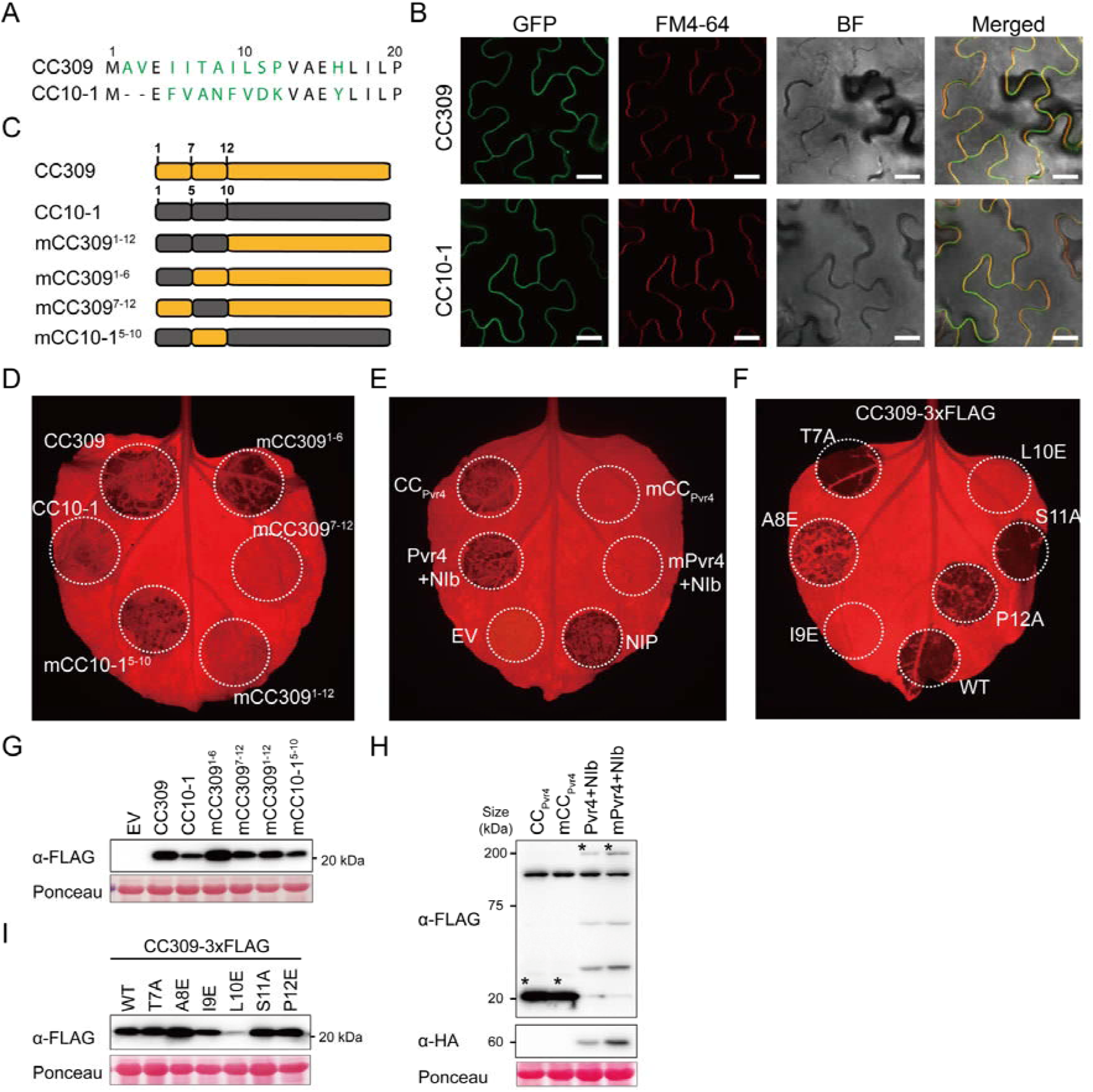
The N-terminal ‘TAILSP’ motif in subclade III G10-CCs is critical for autoactivity. (A) Alignment of the N-terminal region of CC309 and CC10-1. Different residues are highlighted in green. (B) Co-localization of CC309-GFP and CC10-1-GFP with FM 4-64– labeled plasma membrane. Confocal microscopy images were taken at 26 hpi; bar = 20 μm. (C) Schematic diagram of CC309 N-terminal chimeras with CC10-1. (D) Cell death induced by the CC309 and CC10-1 mutants described in (C). mCC10-1^5-10^ (left lower circle) and mCC309^1-6^ (right top circle) contain a ‘TAILSP’ motif. (E) Compromised cell death associated with defective ‘TAILSP’ motif in Pvr4 and the CC domain of Pvr4 (CC_Pvr4_). *Phytophthora sojae* necrosis-inducing protein (NIP) was used as a positive control for cell death. (F) Compromised cell death associated with I9E (left lower circle) and L10E (right top circle) point mutations in the TAILSP motif. All swap- or point mutants were transiently expressed in *N. benthamiana*, and cell death was visualized after 3 dpi. (G-I) Protein accumulation as described in (D), (E), and (F), respectively, was examined by Western blotting analysis. Equal protein loading was confirmed by staining membranes with Ponceau S. The above experiments were repeated at least three times with similar results.

Based on the above observations, we focused on the N-terminus of members of subclade III and identified a well-conserved ‘TAILSP’ motif in members of subclade III, except for CC10-1 (*SI Appendix,* Fig. S4). To confirm the autoactivity function of this motif, we generated CC309 mutants in which 1 to 12 aa or a partial region were substituted with those of CC10-1 (Fig. 3C) and expressed the mutants in *N. benthamiana*. Substitution of residues 7–12 (mCC309^7-12^), corresponding to the ‘TAILSP’ motif, as well as residues 1–12 (mCC309^1-12^), but not residues 1–6 aa (mCC309^1-6^), abolished cell death–inducing activity (Fig. 3D). Conversely, mCC10-1^5-10^ carrying residues 7–12 of CC309 exhibited slightly recovered activity, indicating that residues 7–12 of CC309 are critical for autoactivity. To confirm that the ‘TAILSP’ motif is also critical for R gene-mediated cell death, this motif in CC_Pvr4_ and Pvr4 was mutated to that of CC10-1. Like CC309, mutation of the ‘TAILSP’ motif disrupted the cell death-inducing activity of CC_Pvr4_, even when expressed with its cognate effector, NIb (Fig. 3E). Immunoblotting analysis revealed comparable expression levels of these protein fragments (Fig. 3G, 3H). These results suggest that the ‘TAILSP’ motif of the α1 helix is crucial for autoactivity of G10-CCs and R genes.

Next, to identify the amino acids responsible for functionality, site-directed mutagenesis using synthetic oligonucleotides was performed to exchange each residue in the ‘TAILSP’ motif to an alanine or glutamate. A8, I9, and L10 were substituted with negatively charged glutamate (E), and T7, S11, and 12P were substituted with alanine (A). The A8E mutation substantially reduced the activity, and the I9E and L10E mutations completely disrupted function (Fig. 3F), indicating that hydrophobic residues are essential for cell death-inducing activity. All of the mutant proteins accumulated to similar levels, except for L10E, when expressed in *N. benthamiana* leaves, indicating that the observed loss-of-function phenotype was not due to protein destabilization (Fig. 3I). Collectively, our results suggest that hydrophobic amino acids in the α1 helix of pepper G10-CC are critical for autoactivity.

### Four α helices of G10-CC are required for cell death–inducing activity

It was reported that only 29 amino acids of NRC4 are sufficient to induce cell death in *N. benthamiana* (47). In attempting to define the minimal region of G10-CC sufficient for autoactivity, we found that the G10-CC domain has five α helices via secondary structure analysis. We therefore generated N-terminal or C-terminal deletion mutants of CC309 (Fig. 4A). Truncated fragments of CC309 with a deletion of 5 (N1) or 10 (N2) amino acids at the N-terminus exhibited a diminished cell death-inducing activity (Fig. 4B). C-terminal deletion mutants C1 and C2, but not C3, induced cell death (Fig. 4B and 4D), suggesting that the four α helices of G10-CC are necessary and sufficient for triggering cell death. The capacity for inducing cell death was quantitatively measured based on photosynthetic parameters: the quantum yield of photochemistry in PSII (Fv/Fm) (Fig. 4D) (48). An immunoblot analysis was performed to assess accumulation of the FLAG-fused deletion mutants after 24 h of agroinfiltration. Proteins N1 and N2 accumulated at much lower levels than full-length CC309, suggesting that the N-terminus plays a role in maintaining protein stability. However, although C2 accumulated to a lower level than C1, it exhibited the strongest cell death-inducing activity and lowest Fv/Fm value (Fig. 4B, 4C, and 4D), indicating that protein level does not affect cell death-inducing activity and the α5 helix is not essential for autoactivity. Consistent with previous observations in motif-swap experiments, these results suggest that the α1 helix is critical for the cell death-inducing activity of G10-CC and at least four α helices are required for autoactivity.

**Figure 4.**
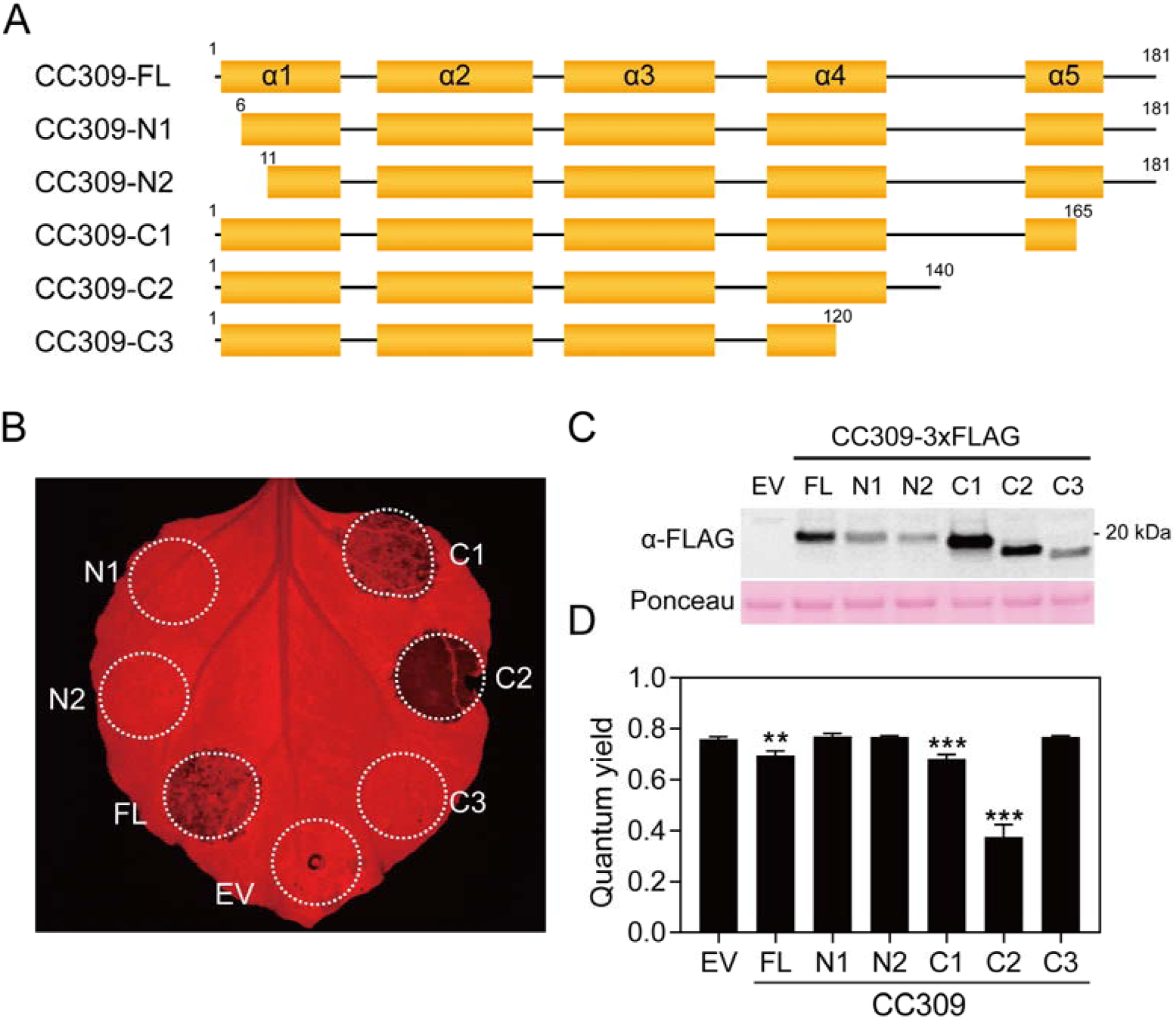
A minimum of four α-helices (α1– α4) are required for the cell death–inducing activity of CC309. (A) Schematic diagram of CC309 and deletion mutants. Secondary structure of CC309 was predicted using JPRED4. Predicted α-helix is indicated by a yellow box. (B) The compromised cell death phenotype associated with the α1 (N1 and N2) or α4 helixes (C3). Deletion mutants were transiently expressed in *N. benthamiana*, and the image was taken at 3 dpi. (C) Protein accumulation was analyzed by immunoblotting. (D) The degree of cell death is reported as quantum yield (Fv/Fm) determined using a closed FluorCam. Significance was determined using one-way ANOVA followed by Dunnett’s multiple comparisons test, with asterisks denoting statistically significant differences: **P < 0.01, ***P < 0.001. Data are the mean (± SD) of four biological replicates.

### Cell death mediated by G10-CCs is independent of helper NLRs

Recent studies revealed that NLR-mediated immunity involves a complex network and many NLRs require helper NLRs to exert activity in the immune response (16, 22). We therefore tested the dependence of G10-NLR- and G10-CC-mediated cell death activity on known helper NLRs. Virus-induced gene silencing (VIGS) was performed to simultaneously co-silence the RNL-type helpers *NbNRG1* and *NbADR1* and the triple NRC-type helpers NRC2, 3, and 4. Autoactive G10-CCs were expressed in silenced plants by agroinfiltration (Fig. 5A). Silencing efficiency was estimated by qRT-PCR analysis of the transcript levels of each gene at 3 weeks after VIGS (Fig. 5C). One of the *NbNRC*-dependent *Phytophthora infestans* resistance genes, *R8*, and its cognate effector, *Avr8*, were used as controls (16). In NRC-silenced plants, cell death mediated by *R8*/*AvrR8* was compromised, whereas cell death mediated by autoactive CC309, NLR620 and Pvr4/NIb was not (Fig. 5A). Cell death induced by G10-CC and G10-NLR was also not compromised in *NRG1-* and *ADR1*-silenced plants (Fig. 5A and 5B). These results indicate that G10-CCs and G10-NLR trigger cell death in a helper-independent manner.

**Figure 5.**
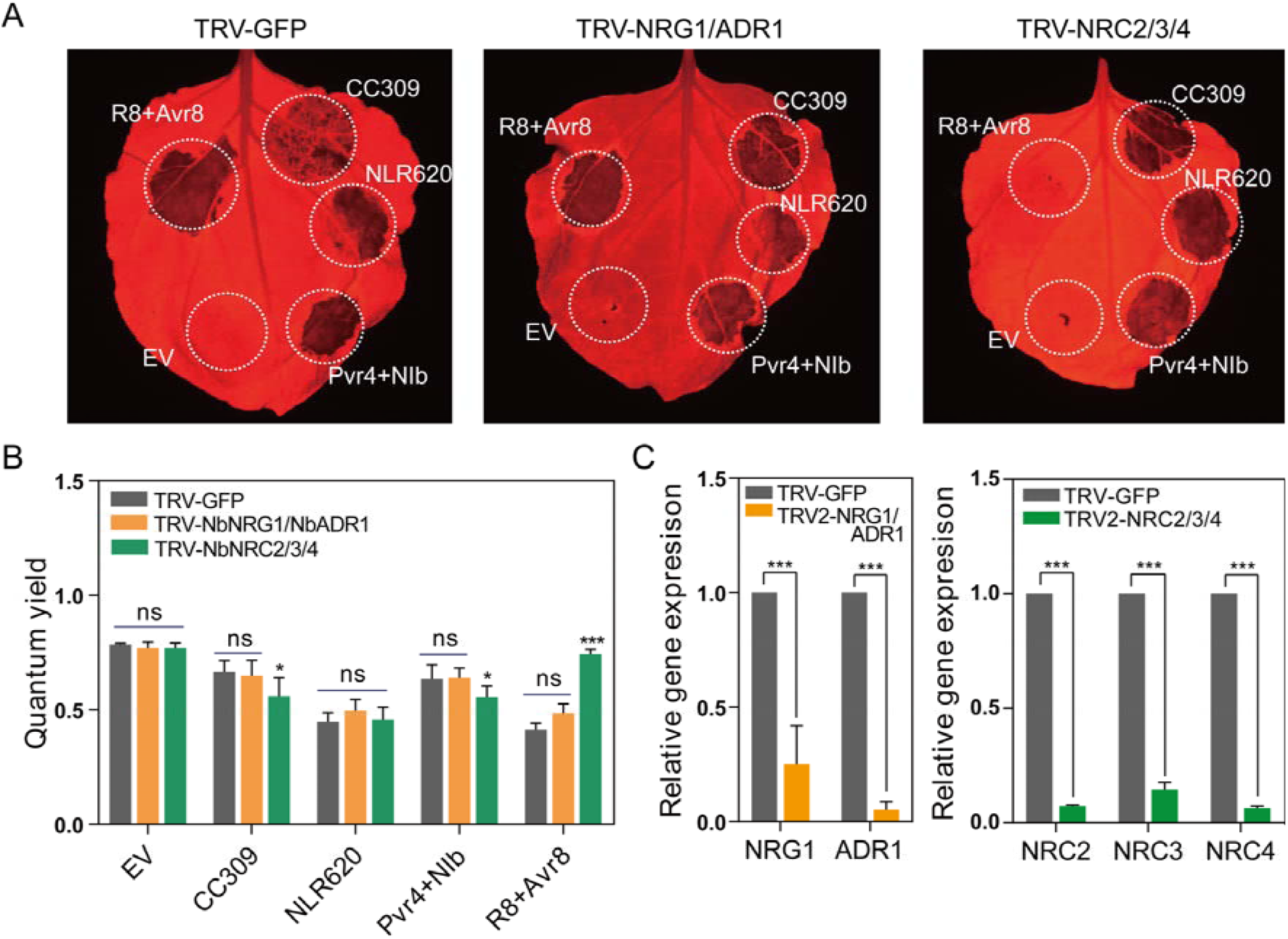
Cell death induced by autoactive G10-CCs or G10-NLRs is independent of NRG1/ADR1*-* or NRC-type helper-mediated pathways. (A) Cell death phenotype induced by autoactive CC309, NLR620 and Pvr4 with NIb in *NRG1*/*ADR1*- or NRC2/3/4-silenced *N. benthamiana*. Agrobacteria carrying each clone were infiltrated 3 weeks after VIGS. The NRC2/3/4-dependent *P. infestans* resistance gene R8 and its cognate effector, Avr8, were used as helper-dependent controls. Each experiment was replicated independently two times (n = 10). Leaves were photographed at 3 dpi. (B) The degree of cell death is reported as quantum yield (Fv/Fm) determined using a closed FluorCam. Significance was determined using one-way ANOVA followed by Dunnett’s multiple comparisons test, with asterisks denoting statistically significant differences: *P < 0.05, ***P < 0.0001. Data are the mean (± SD) of four biological replicates. (C) Relative transcript abundance for each gene was determined by quantitative RT-PCR analysis of silenced plants. The mean values (± SD) for transcript levels were normalized to that of *N. benthamiana EF1-* α. Transcript levels of GFP-silenced plants were set to 1. Error bars represent the mean of three biological replicates, and asterisks denote significant differences at P < 0.0001 as determined by two-way ANOVA followed by Sidak’s multiple comparisons test.

### G10-NLRs are conserved in seed plants, and G10-CCs of other plant species induce cell death

A phylogenetic analysis was conducted on a larger scale to more completely elucidate the divergence and evolution history of NLRs in seed plants. Because it is the only conserved domain of NLR proteins suitable for sequence alignment, the NB-ARC domain of a total 2,419 NLRs from ten representative plant species, including four Solanaceous species (pepper, tomato, potato, and tobacco [*Nicotiana tabacum*]); a Brassicaceae plant (*A. thaliana*); a monocot, Poaceae rice (*Oryza sativa*); a Piperaceae black pepper (*Piper nigrum*), a basal angiosperm (*Amborella trichopoda*); a gymnosperm, Norway spruce (*Picea abies*) and an oldest lineages of vascular plants Selaginella (*Selaginella moellendorffii*) were subjected to phylogenetic analysis. The groups were assigned based on a previous study of the classification of Solanaceae NLRs. Most of the NLRs were included in assigned groups, except for three NLRs of *Selaginella moellendorffii*. G10 formed a monophyletic clade with a high bootstrap value of 91% and was clearly distinguishable from the other CNL clades, despite being typical CNLs (Fig. 6A). We also found that NLRs of nine of ten species (except *S. moellendorffii*) were present in the G10 clade. To determine whether G10 is a unique clade among a wide variety of species, we analyzed the NLR repertoires in each plant species based on their clustering in the phylogenetic tree. Interestingly, G10 was present in all of the plant species examined, even in the most ancestral angiosperm and gymnosperm plants (Fig. 6B, Dataset S3). These results suggest that G10-NLRs are distinct from other conventional CNLs and have existed since the emergence of seed plants.

**Figure 6.**
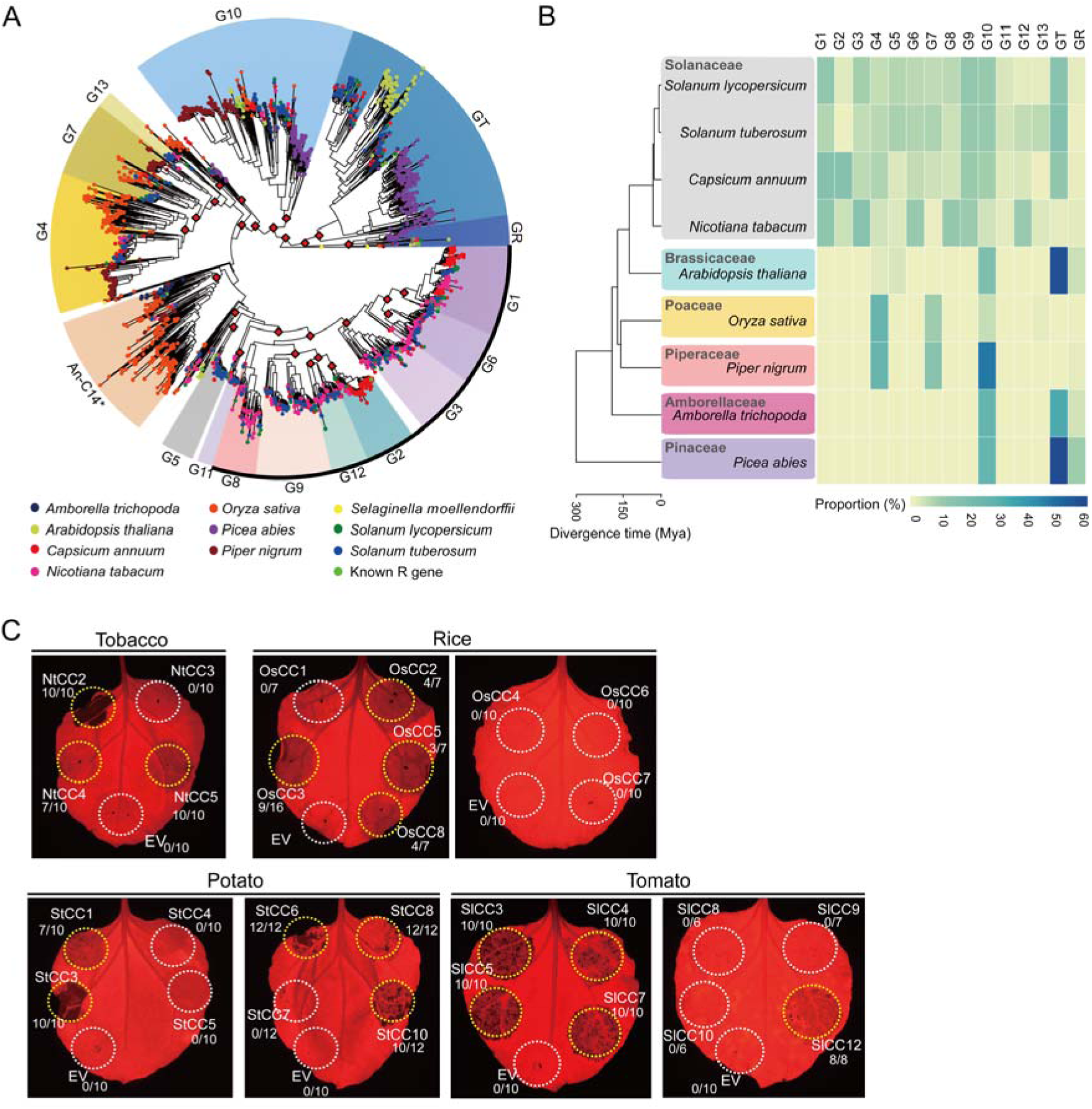
G10-NLRs are conserved across seed plants, and G10-CCs of other plant species also induce cell death. (A) Intact NB-ARC domains in NLRs from 10 plant genomes were used to reconstruct the phylogenetic tree. Groups were assigned based on ultrafast bootstrap (UFBoot) values guided by classification of Solanaceae NLRs. Two NB-ARCs in *S. moellendorffii* were used as the outgroup. UFBoot values >90% are marked as red diamonds at the nodes. Black outline indicates NRC helper-dependent groups. Asterisk denotes the clade described as An-C14 in Shao *et al*. (B) The species tree (left) of nine plant genomes used in this study indicates their evolutionary relationships and divergence time. The proportion of NLRs of each group in plant species was visualized using a heatmap (right). (C) G10-CCs from Solanaceae plants (potato, tomato, and tobacco) and a monocot plant (rice) were expressed in *N. benthamiana*. Autoactive cell death is indicated by yellow circles, and no cell death is indicated by white circles. The frequency of cell death induced by G10-CC is presented below. Images were acquired at 3 dpi. The experiments were repeated three times with at least six biological replicates.

Next, we examined the commonality of G10-CC-medicated cell death among other plants. Representative G10-CCs from five plant species (pepper, tomato, potato, tobacco, and rice) were randomly chosen from a subclade in the phylogenetic tree (Dataset S4). More than 40% of tested G10-CCs from Solanaceae plants and even taxonomically distant rice trigger cell death in *N. benthamiana* (Fig. 6C, *SI Appendix,* Table S3). In addition, a recent study reported that significantly high portion of the predicted CC domain of the CNLs in *A. thaliana* ecotype Columbia-0 triggered cell death (49). All of the CNLs from group B described by Wróblewski *et al*., except At1g63360, belonged to the G10-NLR group according to our classification, and 13 of 22 G10-CC domain fragments induced cell death in *N. benthamiana* (49). Considered in light of this previous report, our results indicate that the G10-CCs of not only Solanaceae plants but also evolutionarily distant rice trigger cell death.

## Discussion

NLR proteins are one of the largest and most widespread families of intracellular immune receptors in plants. These proteins activate potent immune responses that often accompany a HR. In this study, we demonstrated that among the 14 known NLR groups in pepper, G10 is a major NLR with an autoactive NTD. Autoactive G10-CC-induced cell death resembles R gene-mediated HR and immune responses, suggesting that G10-NLRs can trigger immune signaling. Analyses of pepper G10-CC sequences revealed that non-autoactive CCs contain partial deletions or variations in the α1 helix region and the ‘TAILSP’ motif is critical for cell death-inducing activity. G10-CC autoactivity requires a minimum of four α-helices. Furthermore, cell death mediated by G10-CCs is independent of helper NLRs. Remarkably, G10-NLRs are widely conserved in seed plants, and G10-CCs of other plant species, such as Solanaceae and even non-Solanaceae plants, induce cell death in *N. benthamiana*.

Some NLRs function as sensor/signaling pairs. Sensor NLRs contain additional unusual non-conserved domains that function in pathogen recognition and are coupled to a helper NLR (also known as an ‘executor’) to initiate immune signaling (12). Sensor NLRs form heterodimers with executor NLRs to suppress autoactivity in the absence of pathogen detection (13, 14). The recognition of a pathogen by a sensor NLR leads to a change in the executor NLR state from inactive to active. Generally, NLR pairs are genetically linked and oriented head-to-head, sharing a promoter for simultaneous regulation of transcription (15, 50). We found that autoactive G10-NLRs are not oriented head-to-head in the upstream region and do not contain the additional integrated domain (*SI Appendix,* Table S1). Additional genome-wide analyses using updated pseudomolecules identified 19 NLR pairs in a head-to-head orientation in pepper (38). Only one G10-NLR (referred to as CaNBARC403) was identified; however, CaNBARC403 does not exhibit autoactivity and does not have an additional domain in its upstream flanking gene (*SI Appendix,* Table S4). These data suggest that autoactive G10-NLRs in pepper do not function as executors in NLR pairs and their activity is regulated by negative regulatory protein(s) rather than inactive-state NLRs, such as RIN4, which negatively regulates RPS2 (51). These data indicate that conventional paired NLRs do not exist in pepper G10-NLRs.

Based on several lines of evidence, we propose that pepper G10-NLRs should be classified as a new putative singleton NLR clade. First, we found that many G10-CCs from Solanaceae plants, including pepper, potato, tobacco and tomato, trigger autoactivity in *N. benthamiana* (Table1, *SI Appendix,* Table S3 and Fig. 6). In addition, a recent study reported that 13 of 22 *Arabidopsis* CNLs in group B that belong to G10 in our classification trigger cell death in *Arabidopsis* (49). Taken together, these results indicate that many G10-CCs in Solanaceae and Brassicaceae plants have the potential to trigger cell death (*SI Appendix,* Table S3). Second, G10-CCs from taxonomically distant plants triggered visible cell death in *N. benthamiana*. We demonstrated that G10-CCs of monocot rice induce cell death in *N. benthamiana* (Fig. 6, *SI Appendix,* Table S3). In addition, among 22 *Arabidopsis* G10-CCs, 13 in *N. benthamiana* and 5 in lettuce have been shown to induce cell death (49). Third, a functional analysis of known R genes reported that G10-NLRs specifically sense cognate effector proteins (24, 37). The CC domain of the functional G10-NLR RPS5 was shown to play an important role in recognizing the cognate effector AvrPphB (24). Previously, we reported that the LRR domain of Pvr4 is important for recognition of PepMoV-NIb, based on a domain-swap experiment with the susceptible allele of Pvr4 (37). Taken together, these data indicate that G10-NLRs function in effector recognition as well as initiation of immune signaling. Finally, the proposed classification of pepper G10-NLRs as a new putative singleton NLR clade is also supported by the finding of sustainable cell death mediated by G10-CC or G10-NLR in helper NLR-silenced plants such as RNLs and NRCs (Fig. 5).

The phylogenetic tree analysis indicated that G10 is an NRC-independent clade (Fig. 6A). However, it is doubtful that all G10-NLRs across plant species utilize the same mechanism to confer resistance, as the helper NLRs ADR1 and NRG1 are either completely or partially required for cell death mediated by RPS2, a G10-NLR in *Arabidopsis* (22). Interestingly, we found that RPS2 also induces cell death in helper NLR-silenced *N. benthamiana* (unpublished data). Therefore, an alternative scenario is that G10-NLRs do not require other NLR genes to function in evolutionarily distant plants, and downstream signaling components for G10-mediated cell death are well conserved over a wide range of plant species. Adachi *et al.* recently suggested that NLRs evolved from multifunctional singleton receptors into functionally specialized and diversified receptor pairs and intricate receptor networks (8). We demonstrated that G10 forms a distinct phylogenetic cluster that includes several cloned functional R genes: *Pvr4* and *Tsw* in pepper and *RPS2* and *RPS5* in *Arabidopsis* (34, Fig. 6A, Dataset S3). We also demonstrated that G10 is the only one well conserved across multiple plant species, including the basal angiosperm *A. trichopoda* and the gymnosperm *P. abies* (Fig. 6B, *SI Appendix,* Table S5). A previous study also reported that the AN-C2 group corresponding to G10 is an ancestral angiosperm lineage, suggesting that G10-NLRs are ancient compared with other CNL groups (52).

In this study, using an unprecedented high proportion of tested CC domains compared to other groups, we found that pepper G10-CCs exhibit autoactivity (Table 1). However, G10-NLRs are not the only group that contains autoactive CC domains. It was reported that transient overexpression with the tomato or tobacco CC domain of the I2-like NLR induces cell death in *N. benthamiana*, but this does not occur with the CC domain of pepper I2-like NLRs (31). Interestingly, C-terminal tagging of the CC domain of pepper I2-like NLR with enhanced yellow fluorescent protein (note that the pepper I2-like NLRs belong to G4) strongly triggers cell death upon transient overexpression in *N. benthamiana.* This tag-dependent autoactivity has been observed with several CC domains of NLRs, regardless of the level of protein accumulation (31). Both GFP and YFP are known to have a weak tendency toward dimerization, which could have an effect on autoactivity. These results suggest that in pepper G4-NLRs, the CC domain evolved differently after speciation to require additional domain(s) for oligomerization to trigger immune signaling; mechanistically, G4- and G10-NLRa appear to behave differently in terms of inducing cell death.

Analyses of pepper G10-CC sequences indicated the presence of partial deletions or variations in the α1 helix region in non-autoactive CCs (Fig. 2 and *SI Appendix,* Fig. S4). Mutations in the α1 helix of G10-CC or G10-NLR compromised their cell death-inducing activity (Fig. 3). This region is matched with a ZAR1 α1 helix forming funnel-shaped structure in the PM after conformational switching during activation of the ZAR1 resistosome, and amphipathic residues in the ZAR1 α1 helix are known to be essential for cell death-inducing activity (32, 47). Collectively, these data indicate that the α1 helix is important for triggering cell death in singleton NLRs. However, the α1 helix of pepper G10-CCs is required for cell death-inducing activity but does not affect PM localization. It would be interesting to examine in future experiments whether pepper G10-CCs or G10-NLRs form multimers similar to the ZAR1 resistosome in order to elucidate the general mechanism of NLR-induced cell death. Here, we suggest that G10-NLRs not only serve as valuable resources for increasing our understanding of the molecular basis of NLR-mediated cell death but also provide critical information for identifying new disease-resistance genes in crop plants.

## Materials and Methods

### Plant materials and growth conditions

*Nicotiana benthamiana* plants were grown in horticultural bed soil (Biogreen, Seoul Bio Co., Ltd., Seoul, Korea) in a well-maintained chamber under conditions of 16-h/8-h photoperiod, temperature of 25°C, photosynthetic photon flux density of 80∼100 µmol m^−2^ s^−1^ and relative humidity of 70%. Four-week-old plants were used for transient overexpression. Foliage leaves of 2-week-old plants were inoculated with *Agrobacterium* for VIGS.

### Identification of NLRs and phylogenetic tree analysis

Annotated protein sequences of *Capsicum annuum* v1.55 (53), *Nicotiana tabacum* Nitab v4.5 (54), *Oryza sativa* RGAP 7.0 (55), *Picea abies* v1.0 (56), *Piper nigrum* v1.0 (57), *Selaginella moellendorffii* v1.0 (58), *Solanum lycopersicum* ITAG 2.4 (59), *Solanum tuberosum* PGSC 3.4 (60) and known NLR genes from the Genbank and Plant Resistance Genes database v3.0 (61) were used in this study. NLR identification and classification methods were based on a previous study (34), with some modifications. To identify NLRs containing the NB-ARC domain (PF00931), InterProScan v5.22-61 (62) was performed with default parameters. Subsequently, these NLRs were scanned with MAST implemented in MEME v4.9.1 (63) using NB-ARC motif information from NLR-Parser v1.0 (64, 65). NB-ARC domains with at least three of the four major motifs (P-loop, GLPL, Kinase2 and MHDV) in order and a length of at least 160 aa were selected as intact NB-ARC domains and aligned using MAFFT v7.407 (--maxiterate 1000 --globalpair) (66). Positions containing gaps of ≥90% in multiple sequence alignment were trimmed using TrimAl v1.4.rev22 (67). Phylogenetic relationships were reconstructed based on the maximum-likelihood method using IQ-Tree v1.6.12 (68), with 1000 ultrafast bootstrap approximation (UFBoot) (-bb 1000-safe) (69). Substitution models were selected using ModelFinder (70) implemented in IQ-Tree. The best-fit model was JTT+F+R9. NLR groups were assigned based on known NLR genes, UFBoot value >90% and previously assigned group information (34). Phylogenetic trees and heatmaps of the proportion of intact NB-ARC domains for each NLR group in plant species were plotted using ggtree v1.6.11 (71, 72). The plant species tree was generated using TimeTree (73).

### Plasmid construction

Fragments of pepper NLRs were amplified from genomic DNA of *Capsicum annuum* L. cv Criollo de Morelos 334 (CM334) based on the pepper reference annotation v.1.55 (53). To amplify target genes, primers were specifically designed within 1 kb of the 5’- or 3’-UTR for each gene. Amplicons were cloned into the pCAMBIA2300-LIC vector containing a cauliflower mosaic virus 35S promoter and nopaline synthase terminator using the ligation-independent cloning (LIC) method (36, 74). The N-terminal domain of an NLR was defined as spanning from methionine to just before the p-loop motif in the NB-ARC domain. Amplified NTD fragments were cloned into the pCAMBIA2300-LIC or pCAMBIA2300-3xFLAG-LIC vector in the same manner as full-length NLRs. For cloning of CC domains from other plant species, the fragments were amplified from genomic DNA of rice (*Oryza sativa* L. cv. Nakdong) and tobacco (*Nicotiana tabacum* cv. Xanthi) or cDNA of tomato (*Solanum lycopersicum* cv. Heinz) and potato (*Solanum tuberosum*). To silence *NRC2/3/4*, 285 bp of *NbNRC2*, 334 bp of *NbNRC3* and 272 bp of *NbNRC4* were combined by overlap PCR, and then the overlapped fragment was cloned into the pTRV2-LIC vector. For RNL silencing, a 283-bp fragment of *NbNRG1* and 329-bp fragment of *NbADR1* were combined by overlap PCR, and then the overlapped fragment was cloned into the TRV2-LIC vector. The primers used for PCR in this study are listed in Dataset S4.

### Site-directed and motif-swap mutagenesis

N-terminal motif-swap and site-directed mutants were generated using specific primers carrying the desired mutations. The amplified fragments for tag fusion proteins were cloned into the pCAMBIA2300-3xFLAG-LIC or pCAMBIA2300-eGFP-LIC vector using a ligation-independent cloning method.

### Disease-resistance assay

For PVX:GFP co-expression, *A. tumefaciens* GV3101 carrying the PVX:GFP-expressing binary vector was co-infiltrated with G10-CCs, G10-NLRs and Pvr4/NIb. At 30 hours post-infiltration (hpi), the intensity of GFP fluorescence was measured using a closed FluorCam (Photon Systems Instruments, Czech Republic) with a GFP filter to quantify PVX replication.

### *Agrobacterium*-mediated transient overexpression in *N. benthamiana*

*Agrobacterium tumefaciens* GV3101 strains carrying the various constructs were prepared for transient overexpression. Bacteria were grown overnight at 28°C in LB medium supplemented with kanamycin (50 μg/ml) and rifampicin (50 μg/ml). The cells were then pelleted and resuspended in infiltration buffer (10 mM MES [pH 5.6] and 10 mM MgCl_2_ with 150 μM acetosyringone) at an optical density at 600 nm (OD_600_) of 0.3 for CC domains or 0.6 for full-length NLRs and R genes. To screen G10-CCs from tomato, tobacco, potato, and rice, G10-CCs were co-expressed with the gene-silencing suppressor, p19 (final OD_600_ = 0.25). Agrobacterial suspensions were applied to infiltrate the abaxial leaves of 4-week-old *N. benthamiana* plants using a needleless syringe. Macroscopic cell death phenotypes were detected using a fluorescence-labeled organism bioimaging instrument system (Neoscience, Korea). The degree of cell death was reported as quantum yield (Fv/Fm) using a closed FluorCam (Photon Systems Instruments).

### Virus-induced gene silencing

VIGS was performed in *N. benthamiana* as described by Liu *et al.* (75). Suspensions of pTRV1 and pTRV2 carrying the target gene fragment were mixed in a 1:1 ratio in infiltration buffer at a final OD_600_ of 0.15. Two leaves of 2-week-old *N. benthamiana* seedlings were infiltrated with the *Agrobacterium* suspension mixture. Three weeks later, the upper leaves were collected to confirm silencing of the target gene and used for subsequent experiments.

### Gene expression assay

Total RNA was extracted using TRIzol reagent (MRC, OH, USA), and cDNA was synthesized using Superscript II (Invitrogen, CA, USA). Gene-specific primers were used for quantitative RT-PCR at 95°C for 5 min, followed by 40 cycles of denaturation at 95°C for 15 s and 55°C for 1 min. qRT-PCR was performed using a CFX96 real-time PCR instrument (Bio-Rad, USA) with SsoAdvanced universal SYBR green supermix (Bio-Rad). Nucleotide sequences of all primers used in this study are listed in Dataset S3. Gene transcript levels were normalized to that of the elongation factor gene, *NbEF1-* α.

### Confocal laser scanning microscopy

Cells expressing GFP-tagged proteins were examined using a Leica confocal microscope SP8X (Leica Microsystem, Germany) with 40×/1.0 water-dipping objective. GFP-expressing cells were imaged using 488-nm excitation and detection of emission from 500–530 nm. Simultaneous excitation of GFP and plasma membrane marker dye, FM4-64 (Invitrogen), was performed using 488-nm excitation and detection of emission signals at 500–530 nm and 600–650 nm for GFP and FM4-64, respectively.

### DAB staining

Accumulation of H_2_O_2_ was monitored by DAB staining. *Nicotiana benthamiana* leaves were detached and incubated in DAB-HCl solution (1 mg/ml, pH 3.8) overnight at 25°C in the dark. After staining, the leaves were soaked in 95% ethanol to remove chlorophyll.

### Immunoblot analysis

Leaf tissue (50 mg) of *N. benthamiana* was ground into fine powder in liquid nitrogen and homogenized in 200 μl of extraction buffer (50 mM Tris-HCl [pH 7.5], 100 mM NaCl, 1 mM EDTA, 1% [v/v] IGEPAL CA-630, 10% glycerol, 5 mM DTT, 1 mM NaF, 1 mM PMSF, 1× EDTA-free protease inhibitor cocktail [Roche]). The lysates were cleared by centrifugation at 12,000 *g* for 10 min at 4°C. Samples were loaded on an 8 or 10% SDS-PAGE gel, and proteins were detected after blotting with anti-FLAG M2 antibody (SIGMA) (1:10,000 dilution) or anti-HA antibody (Abcam) (1:5,000 dilution).

### Quantification and statistical analyses

Statistical analyses and graph generation were performed using Prism7 software (GraphPad). Error bars in all figures represent the standard deviation of the mean. The number of replicates is reported in the figure legends. Statistical comparisons between different samples was carried out by one-way or two-way analysis of variance (ANOVA) with Sidak’s multiple comparisons or Dunnett’s multiple comparisons tests. Samples exhibiting statistically significant differences are marked in the figures with asterisks or letters.

## Acknowledgments

We are thankful to Dr. Hyun-Ah Lee, Dr. Eunyoung Seo, Jihyun Kim and Haeun Kim for construction of NLRs and NTDs. This work was supported by the National Research Foundation of Korea (NRF) grant funded by the Korea government (MSIT) (No. 2018R1A5A1023599 (SRC) and 2018R1A2A1A05019892).

