## Supplementary material for "Genome-wide functional analysis of hot pepper immune receptors reveals an autonomous NLR cluster in seed plants": supplimentary figures

Doil Choi

**This PDF file includes:**

Figures S1 to S8

Tables S1 to S5

**Other supplementary materials for this manuscript include the following:**

Datasets S1 to S4

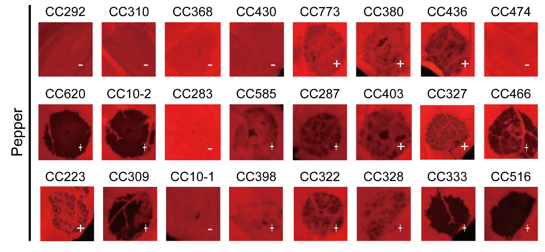

**Fig. S1. Cell death induced by pepper G10-CCs in *N. benthamiana*.**

24 CC domains of pepper G10-NLRs were transiently overexpressed in *N. benthamiana*. Images were taken at 3 days after infiltration. Cell death and no visible cell death are presented + and - , respectively. Experiments were repeated three times.

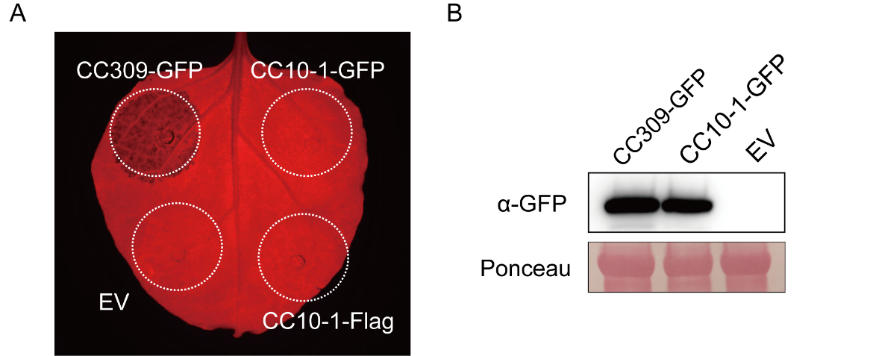

**Fig. S2. Cell death phenotype of GFP-tagged CC309 and CC10-1.**

(A) CC309-GFP and CC10-1-GFP were expressed in *N. benthamiana* leaves by Agroinfiltration with an empty vector (EV) as a negative control. Images were taken 2 days post infiltration. (B) Protein accumulation was examined by western blot analysis. Equal protein loading was confirmed by membrane staining by ponceau solution.

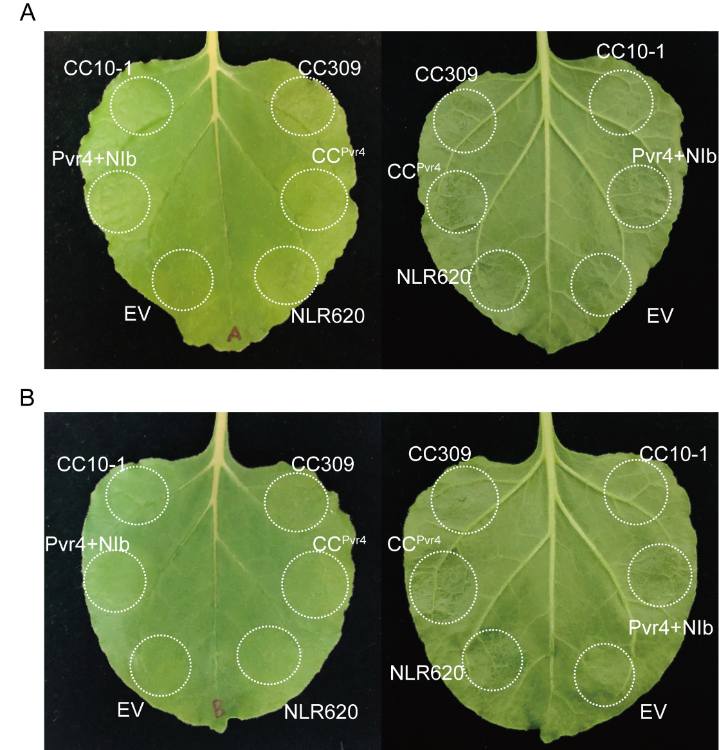

**Fig. S3. No visible cell death in the leaves expressing G10-CCs and G10-NLRs with pCAMBIA2300:GFP or PVX:GFP.**

Leaves were infiltrated pCAMBIA2300:GFP (A) or PVX:GFP (B) with G10-NLR or G10-CCs. Right panel represents adaxial side and left panel shows abaxial side. Same leaves used in Figure 1E were photographed under white light condition 30 hr after agroinfiltration. n=8.

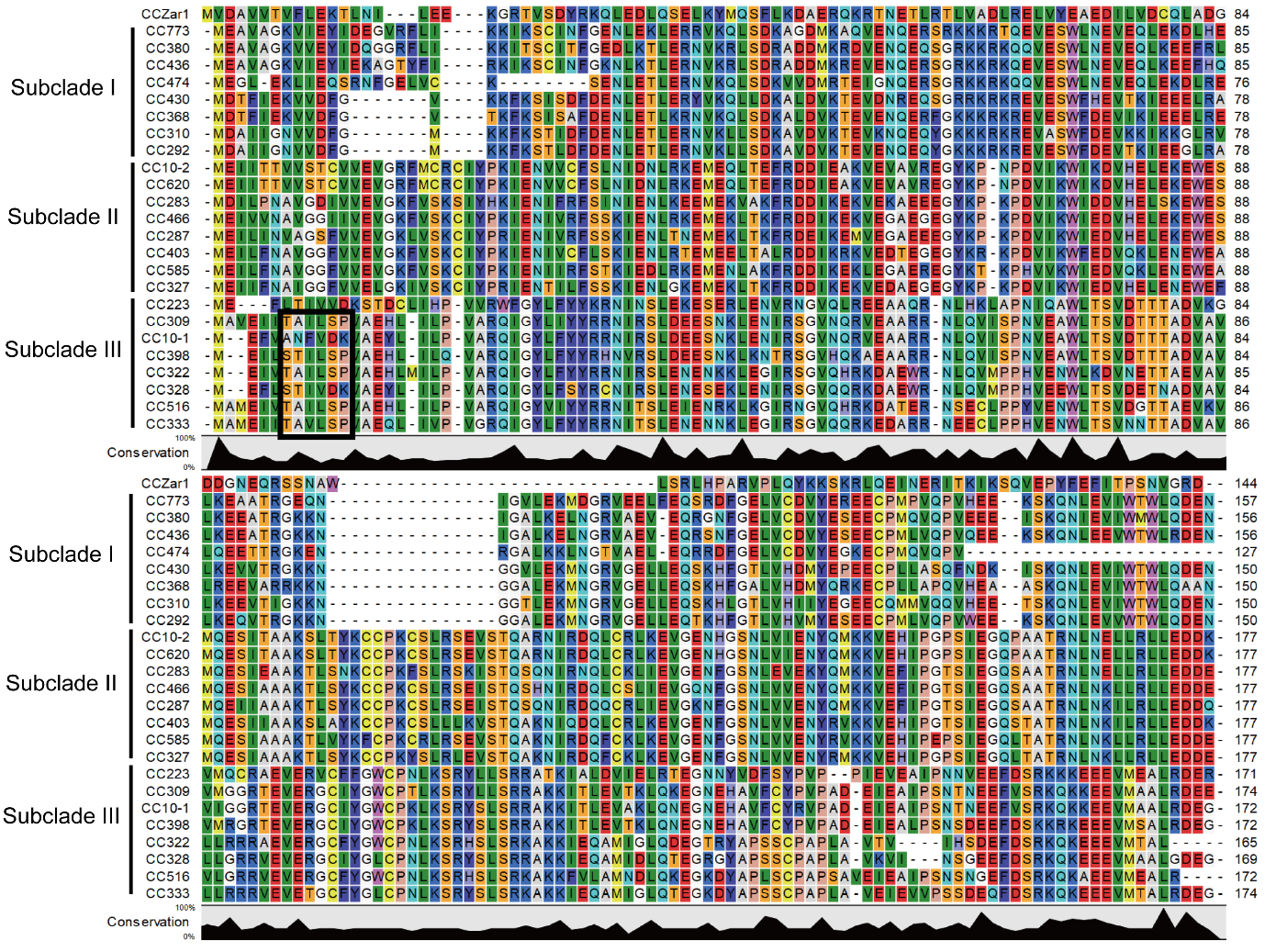

**Fig. S4. Alignment of amino acid sequences of pepper G10-CCs.**

Amino acid sequences of 24 pepper G10-CCs and CC domain of *Arabidopsis* ZAR1 were used for sequence alignment. Alignment were conducted by CLC workbench software. The conserved motif (‘TAILSP’) in clade III was presented in a black box.

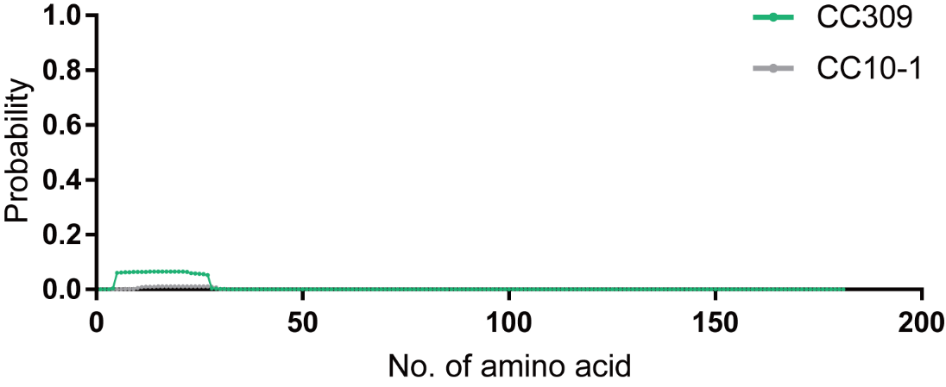

**Fig. S5. Prediction of transmembrane in CC309 and CC10-1 using TMHMM Server v. 2.0.**

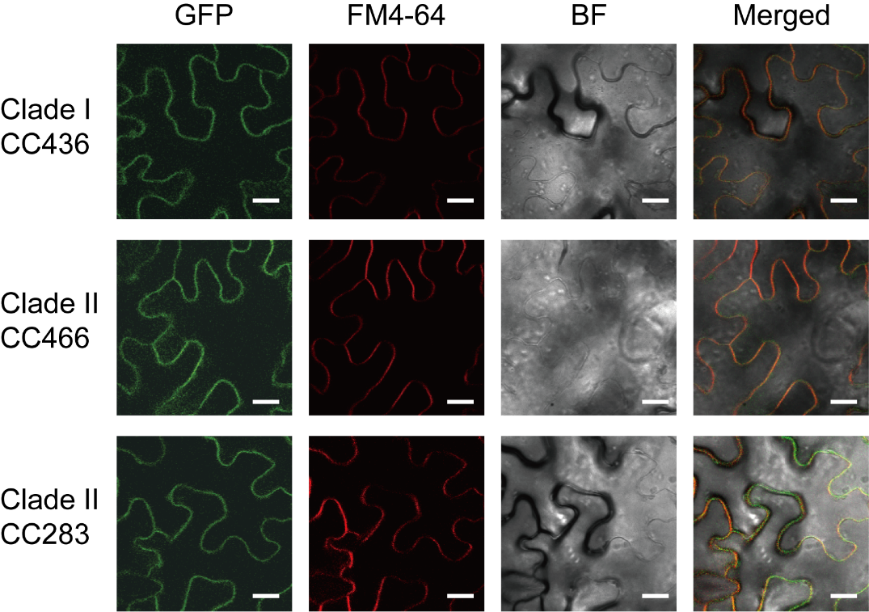

**Fig. S6. Non-autoactive G10-CCs also localize at plasma membrane regardless of their cell death inducing activity**

CC436, CC466 and CC283, one of non-autoactive G10-CCs in subclade I, II, and III, respectively were chosen to explore their subcellular localization. GFP was fused to these proteins at C-terminus and GFP-fused proteins were expressed in *N. benthamiana*. GFP signal was observed by confocal microscopy at 48 hr after infiltration. FM 4-64 dye was used as a PM-marker fluorescence dye. Bar=20 μm.

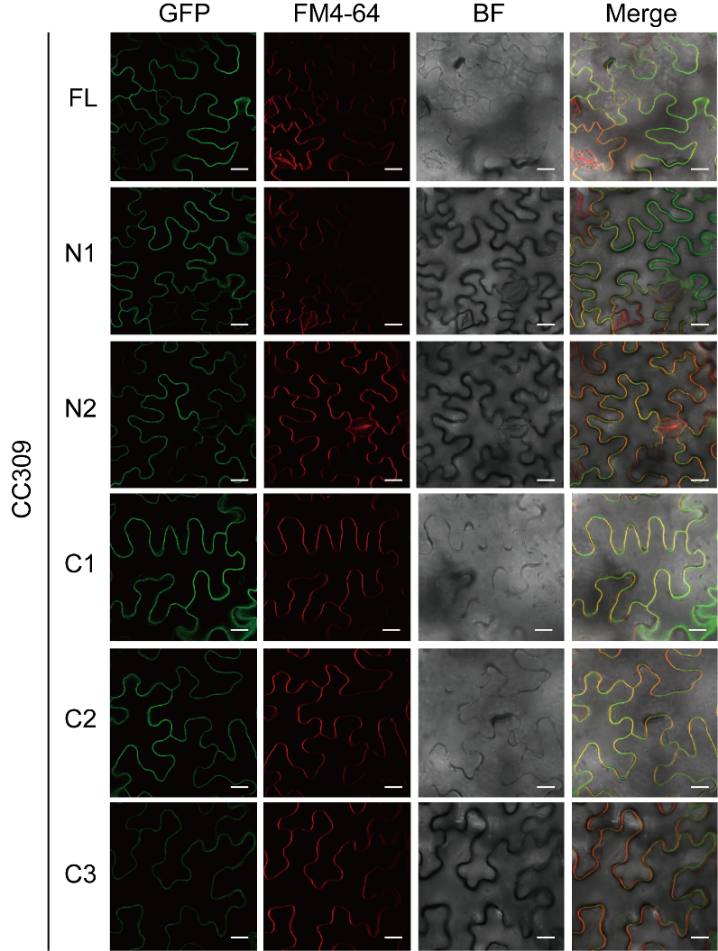

**Fig. S7. The N-terminal or C-terminal deletion mutants of CC309 also localized in the plasma membrane.**

The GFP-fused deletion mutants of CC309 used in Fig. 4 were expressed in *N. benthamiana*. FM 4-64 dye was used as a PM-marker fluorescence dye. Confocal microscopy images were taken at 26 hpi. Bar=20 μm.

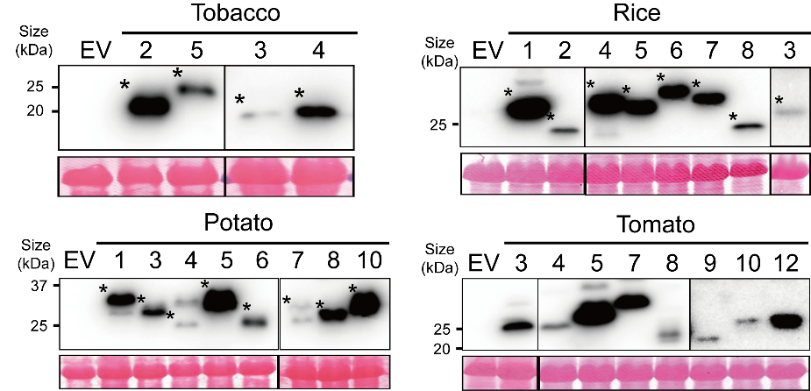

**Fig. S8. The protein expression level of G10-CCs from tobacco, rice, potato and tomato.**

Protein accumulation were analyzed by immunoblot. Asterisks indicate the expected band of proteins. Equal protein loading was confirmed by Ponceau S stained membrane.

**Table S1.** Gene orientation and predicted function of an upstream gene of an autoactive pepper NLR

| Autoactive NLR |  |  | Upstream gene | | |
| --- | --- | --- | --- | --- | --- |
| Gene ID (NLR ID) | Strand |  | Gene ID | Strand | Gene description |
| CA11g06220 (NLR502) | + |  | CA11g06230 | + | Partial TNL-type protein |
| CA12g06200 (NLR504) | + |  | CA12g06190 | - | Putative nuclease HARBI1-like |
| CA12g16200 (NLR582) | + |  | CA00g70880 | - | Ribosomal protein S11 |
| CA12g19860 (NLR530) | + |  | CA12g19850 | + | Partial TNL-type protein |
| CA12g19870 (NLRT2) | + |  | CA12g19860 | + | NLR530 (Autoactive TNL) |
| CA12g19890 (NLRT1) | + |  | CA12g19880 | + | Partial TNL-type protein |
| CA05g05510 (NLR110) | + |  | CA05g05520 | - | Putative LRR receptor-like serine/threonine-protein kinase |
| CA05g02490 (NLR5) | + |  | CA05g02480 | + | Partial CNL-type protein (CNL-NG) |
| CA01g29850 (NLR287) | + |  | CA01g29860 | + | CNL-type protein (CNL-NG) |
| CA01g33790 (NLR620) | + |  | CA01g33800 | + | CNL-type protein (G10) |
| CA01g33850 (NLR10-2) | + |  | CA01g33840 | + | CNL-type protein (G10) |
| CA07g09720 (NLR174) | + |  | CA07g09730 | + | Pentatricopeptide repeat-containing protein |
| CA07g12630 (NLR168) | + |  | CA07g12640 | + | CNL-type protein (CNL-NG) |
| CA08g02000 (NLR95) | + |  | CA08g01990 | + | Partial NL-type protein |
| CA12g19770 (NLR179) | + |  | CA12g19760 | + | Phloem protein 2-like protein |

**Table S2.** List of autoactive NLRs and autoactive N-terminal domains in pepper

| Group | Full-length NLR (NLR ID) | N-terminal domain (NTD ID) |
| --- | --- | --- |
| TNL | CA11g06220 (NLR502) | CA12g19860 (TIR530)* |
|  | CA12g06200 (NLR504) | CA12g19870 (TIRT2)* |
|  | CA12g16200 (NLR582) |  |
|  | CA12g19860 (NLR530)* |  |
|  | CA12g19870 (NLRT2)* |  |
|  | CA12g19890 (NLRT1) |  |
| CNL-G5 | CA05g05510 (NLR110) |  |
| CNL-G9 | CA05g02490 (NLR5) |  |
| CNL-G10 | CA01g29850 (NLR287)* | CA00g76890 (CC403) |
|  | CA01g33790 (NLR620)* | CA01g29850 (CC287)* |
|  | CA01g33850 (NLR10-2)* | CA01g33790 (CC620)* |
|  |  | CA01g33850 (CC10-2)* |
|  |  | CA01g33880 (CC327) |
|  |  | CA01g33900 (CC466) |
|  |  | CA04g10720 (CC585) |
|  |  | CA09g17050 (CC380) |
|  |  | CA09g17080 (CC436) |
|  |  | CA09g17090 (CC773) |
|  |  | CA10g20540 (CC328) |
|  |  | CA10g20630 (CC333) |
|  |  | CA10g20820 (CC223) |
|  |  | CA10g21090 (CC398) |
|  |  | CA10g21120 (CC309) |
|  |  | CA10g21170 (CC322) |
|  |  | CA10g21380 (CC516) |
| CNL-G11 | CA07g09720 (NLR174) | CA03g00800 (CC281) |
| CNL-NG | CA07g12630 (NLR168) | CA02g25800 (RPW595) |
|  | CA08g02000 (NLR95) | CA04g19370 (RPW405) |
|  | CA12g19770 (NLR179) |  |

*: The NLRs that both full length NLR and their N-terminal domain show autoactivity

**Table S3.** The G10-CC domains of other Solanaceae plants as well as rice and *Arabidopsis* are also capable of inducing cell death in *N. benthamiana*.

|  | Assigned G10-NLR | Tested  G10-CC domain | Autoactive  G10-CC domain (%) |
| --- | --- | --- | --- |
| Pepper | 34 | 24 | 17 (70.8) |
| Potato | 32 | 8 | 5 (62.5) |
| Tomato | 15 | 8 | 5 (62.5) |
| Tobacco | 8 | 4 | 3 (75) |
| Rice | 13 | 8 | 4 (50) |
| Arabidopsis* | 23 | 22 | 13 (59.1) |
| * This result is quote from Wro´blewski et al., (2019). | | | |

**Table S4.** Identification of head-to-head oriented NLR genes in pepper genome.

| Gene ID | Type* | Group† | Autoactivity‡ | Integrated  domain | Upstream gene | | | | |
| --- | --- | --- | --- | --- | --- | --- | --- | --- | --- |
|  |  |  |  |  | Gene ID | Type* | Group† | Autoactivity‡ | Integrated  domain |
| CA.PGAv.1.6.scaffold137.47 | N | N.A. | N.D |  | CA.PGAv.1.6.scaffold137.46 | N | N.A. | N.D |  |
| CA.PGAv.1.6.scaffold282.124 | CNL | G9 | N |  | CA.PGAv.1.6.scaffold282.123 | CNL | G9 | N |  |
| CA.PGAv.1.6.scaffold309.40 | CNL | G4 | N |  | CA.PGAv.1.6.scaffold309.39 | CNL | G4 | N |  |
| CA.PGAv.1.6.scaffold551.100 | CN | N.A. | N.D |  | CA.PGAv.1.6.scaffold551.99 | CNL | G2 | N |  |
| CA.PGAv.1.6.scaffold676.64 | CNL | G7 | N |  | CA.PGAv.1.6.scaffold676.63 | N | N.A. | N.D |  |
| CA.PGAv.1.6.scaffold767.32 | CNL | G3 | N | SD | CA.PGAv.1.6.scaffold767.31 | CNL | G3 | N | SD |
| CA.PGAv.1.6.scaffold1084.23 | XN | N.A. | N.D | NACK_C | CA.PGAv.1.6.scaffold1084.22 | CNL | G2 | N |  |
| CA.PGAv.1.6.scaffold1090.4 | CNL | G2 | N |  | CA.PGAv.1.6.scaffold1090.3 | CNL | G2 | N |  |
| CA.PGAv.1.6.scaffold1090.13 | N | N.A. | N |  | CA.PGAv.1.6.scaffold1090.11 | CNL | G2 | N |  |
| CA.PGAv.1.6.scaffold1090.34 | CNL | G12 | N |  | CA.PGAv.1.6.scaffold1090.33 | CNL | G2 | N |  |
| CA.PGAv.1.6.scaffold1090.37 | XN | N.A. | N.D | Haem-oxygenase | CA.PGAv.1.6.scaffold1090.36 | N | N.A. | N.D |  |
| CA.PGAv.1.6.scaffold1357.10 | CNL | G2 | N |  | CA.PGAv.1.6.scaffold1357.9 | CNL | G2 | N |  |
| CA.PGAv.1.6.scaffold1357.16 | CNL | G2 | N |  | CA.PGAv.1.6.scaffold1357.15 | CNL | G2 | N |  |
| CA.PGAv.1.6.scaffold1373.15 | CNL | G2 | N.D |  | CA.PGAv.1.6.scaffold1373.14 | N | N.A. | N.D |  |
| CA.PGAv.1.6.scaffold1578.13 | CNL | G2 | N |  | CA.PGAv.1.6.scaffold1578.12 | CNL | G2 | N |  |
| CA.PGAv.1.6.scaffold1640.8 | CN | N.A. | N.D |  | CA.PGAv.1.6.scaffold1640.7 | CNL | G2 | N |  |
| CA.PGAv.1.6.scaffold1646.8 | CNL | G10 | N |  | CA.PGAv.1.6.scaffold1646.7 | CNL | G1 | N |  |
| CA.PGAv.1.6.scaffold1677.2 | N | N.A. | N.D |  | CA.PGAv.1.6.scaffold1677.1 | XN | N.A. | N.D | NACK_C |
| CA.PGAv.1.6.scaffold1982.2 | NL | N.A. | N.D |  | CA.PGAv.1.6.scaffold1982.1 | N | N.A. | N.D |  |

*NLR type: CNL, CC- *NBARC-LRR; N, NBARC; CN, CC-NBARC; NL, NBARC-LRR; XN, extra domain-NBARC

†Group: N.A., Not assigned; N, No autoactivity

‡Autoactivity: N.D, Not determined; N, No autoactivity

**Table S5.** The number of NLRs for each group in ten plant genome.

|  | G1 | G2 | G3 | G4 | G5 | G6 | G7 | G8 | G9 | G10 | G11 | G12 | G13 | GT | GR | NG | Total |
| --- | --- | --- | --- | --- | --- | --- | --- | --- | --- | --- | --- | --- | --- | --- | --- | --- | --- |
| *Solanum lycopersicum* | 21 | 3 | 14 | 8 | 11 | 10 | 9 | 10 | 20 | 18 | 4 | 2 | 2 | 25 | 2 | 12 | 171 |
| *Solanum tuberosum* | 20 | 0 | 15 | 18 | 23 | 21 | 22 | 12 | 33 | 32 | 9 | 15 | 17 | 52 | 1 | 14 | 304 |
| *Capsicum annuum* | 67 | 69 | 23 | 29 | 9 | 20 | 16 | 13 | 41 | 35 | 7 | 10 | 1 | 52 | 3 | 23 | 418 |
| *Nicotiana tabacum* | 48 | 15 | 56 | 12 | 8 | 40 | 2 | 42 | 36 | 13 | 5 | 38 | 7 | 4 | 5 | 39 | 370 |
| *Arabidopsis thaliana* | 0 | 0 | 0 | 2 | 3 | 0 | 0 | 0 | 0 | 23 | 0 | 0 | 0 | 73 | 5 | 18 | 124 |
| *Oryza sativa* | 0 | 0 | 0 | 98 | 0 | 0 | 34 | 0 | 0 | 17 | 0 | 0 | 3 | 0 | 1 | 222 | 375 |
| *Piper nigrum* | 0 | 0 | 0 | 80 | 0 | 0 | 49 | 0 | 0 | 150 | 0 | 0 | 0 | 0 | 3 | 26 | 308 |
| *Amborella trichopoda* | 0 | 0 | 0 | 0 | 0 | 0 | 0 | 0 | 0 | 17 | 0 | 0 | 0 | 21 | 2 | 23 | 63 |
| *Picea abies* | 0 | 0 | 0 | 0 | 0 | 0 | 0 | 0 | 0 | 92 | 0 | 0 | 0 | 208 | 31 | 15 | 346 |
| *Selaginella moellendorffii* | 0 | 0 | 0 | 0 | 0 | 0 | 0 | 0 | 0 | 0 | 0 | 0 | 0 | 0 | 0 | 3 | 3 |
